# Alzheimer’s environmental and genetic risk scores are differentially associated with ‘g’ and δ

**DOI:** 10.1101/279018

**Authors:** Shea J. Andrews, G. Peggy McFall, Roger A. Dixon, Nicolas Cherbuin, Ranmalee Eramudugolla, Kaarin J Anstey

**Author notes:** Correspondence to: Shea Andrews, Icahn School of Medicine at Mount Sinai 1425 Madison Ave New York, NY, 10029.

## Abstract

**Introduction:** We investigated the association of the Australian National University Alzheimer’s Disease Risk Index (ANU-ADRI) and an AD genetic risk score (GRS) with cognitive performance.

**Methods:** The ANU-ADRI (composed of 11 risk factors for AD) and GRS (composed of 25 AD risk loci) were computed in 1,061 community-dwelling older adults. Participants were assessed on 11 cognitive tests and activities of daily living. Structural equation modelling was used to evaluate the association of the ANU-ADRI and GRS with: 1) general cognitive ability (g) 2) dementia related variance in cognitive performance (δ) and 3) verbal ability, episodic memory, executive function and processing speed.

**Results:** A worse ANU-ADRI score was associated with poorer performance in ‘g’, δ, and each cognitive domain. A worse GRS was associated with poorer performance in δ and episodic memory.

**Discussion:** The ANU-ADRI was broadly associated with worse cognitive performance, validating its further use in early dementia risk assessment.

**Highlights:**

- An environmental/lifestyle dementia risk index is broadly associated with cognitive performance
- An Alzheimer’s genetic risk score is associated with dementia severity and episodic memory
- The environmental risk index is more strongly associated with dementia severity than genetic risk

**Research in Context:** *Systematic Review:* The authors reviewed the literature using online databases (e.g. PubMed). Previous research has highlighted the need for dementia risk assessment tools to be evaluated on outcomes prior to dementia onset, such as cognitive performance. The relevant citations have been appropriately cited.

*Interpretation:* The Australian National University Alzheimer’s Disease Risk Index (ANU-ADRI) was more broadly associated with cognitive performance than Alzheimer’s genetic risk. For the ANU-ADRI, stronger effects were observed for dementia-related variance in cognitive task performance that for variance in general cognitive function. This suggests that ANU-ADRI is more specifically associated with dementia-related processes and further validates its use in early risk assessment for dementia.

*Future Directions:* Accordingly, future studies should seek to evaluate the association of the ANU-ADRI and genetic risk with AD biomarkers and longitudinal cognitive performance to evaluate differential trajectories in ‘g’ and δ.

## 1 Introduction

Given both the projected increase in the prevalence of Alzheimer’s disease (AD) and other dementias and the lack of disease-modifying treatments for AD, risk reduction strategies focusing on prevention to reduce dementia prevalence are required [1,2]. Risk assessment tools are a key component of dementia prevention, as they can provide clear and effective communication of key risk factors that can inform personalized prevention regimes to guide the implementation of risk reduction strategies. As AD has a long preclinical phase, risk assessment tools that can identify at-risk individuals in the early stages of the pathological processes may have significant utility in prevention. As some cognitive decline in normal populations may signal increased risk of dementia, it is important to evaluate and compare the sensitivity of AD risk assessment tools for detecting early dementia-related cognitive disturbance.

Cognitive performance is a combination of an individual’s inherent cognitive ability, decline promoted by the gradual accumulation of neuropathology associated with various chronic conditions of aging [3,4] and cognitive reserve promoting resilience to the pathological changes associated with neurodegeneration [5]. Individual differences in cognitive level and trajectories of cognitive performance with aging are substantial [6]. Nevertheless, performance across multiple tests of cognitive abilities has been observed to be positively inter-correlated. This observation gave rise to the concept of a general factor of cognitive ability (‘g’) representing the shared variance across observed performance on cognitive tasks [7]. Additionally, cognitive tasks that draw on more similar processes tend to be more highly correlated with each other, which can be accounted for by developing additional factors that define specific cognitive domains [8]. A recently proposed extension of ‘g’ for the early detection of dementia is a model that distinguishes dementia-related variance in cognitive task performance (δ) from variance that is unrelated to dementia processes (*g*’) [9,10]. Both *g*’ and δ are derived from a theory-driven confirmatory factor analysis in a structural equation model framework that combines cognitive and functional measures. Functional status is typically measured by activities of daily living (ADL) that represent capacities that are required for autonomous function within society and at home [11]. Given that deficits in cognition and functional status are key characteristics for a clinical diagnosis of dementia, ‘δ’ has been proposed as a measure to detect early cognitive change and concomitant functional decline [9]. For example, δ is related to dementia status (AUC = 0.942) [10], dementia severity (*r* = 0.84) [10], post-mortem AD neuropathology [12], abnormal CSF amyloid-β/Tau biomarkers ratios (AUC = 0.78) [13], cognitive decline and future dementia severity [9,14,15], conversion from MCI to AD (AUC = 0.84) [13], and conversion from normal cognition to MCI or AD (OR = 1.52) [16].

There has been increasing interest in evaluating the effect of AD risk factors with preclinical cognitive performance. For example, genetic risk scores (GRS) composed of the top hits from genome-wide association studies of AD have been associated with faster decline in episodic memory [17,18] and processing speed [18]. Modifiable lifestyle, medical, and environmental risk factors appear to moderate genetic risk of AD and may also have direct effects on vascular cognitive impairment and brain ageing. It has been estimated that 28.2% - 48.4% of dementia cases can be attributed to up to nine modifiable risk factors; specifically, including education, midlife hypertension, midlife obesity, hearing loss, late-life depression, diabetes, physical inactivity, smoking, and social isolation [19-21]. The commonality between these dementia risk factors and those for cognitive decline suggest that risk assessment tools for dementia should also be associated with cognitive decline and cognitive deficits during the preclinical stages [22]. The Cardiovascular Risk Factors, Aging and Dementia (CAIDE) risk score was the first published risk assessment tool for estimating dementia risk and includes midlife measures of vascular risk [23], with higher scores also been associated with faster rates of cognitive decline [23,24]. However, CAIDE was developed for use in midlife cohorts and it is possible the weights developed for CAIDE are study specific [24].

The Australian National University Alzheimer’s Disease Risk Index (ANU-ADRI) is a self-report questionnaire-based risk assessment tool composed of 11 risk and 4 protective factors that were identified using an evidence-based medicine approach [25]. The ANU-ADRI has been validated in three independent cohorts and compared to the CAIDE risk score, where it was found to be predictive of incident AD and all cause dementia [24]. Additionally, the ANU-ADRI was associated with an increased risk of progressing from normal cognition to mild cognitive impairment (MCI) [26] and was found to predict lower brain volumes in the default mode network [27]. However, to date, the association of the ANU-ADRI with cognitive performance has not been evaluated.

The aim of the present study was to expand on this body of research by evaluating the association of the ANU-ADRI in conjunction with an AD GRS with cross-sectional cognitive performance. Cognitive performance was assessed using a comprehensive cognitive test battery in a large community-based cohort of 1,061 older adults without dementia. Three models of cognition were constructed using confirmatory factor analysis (CFA) representing:1) general cognitive ability (g); 2) dementia-related variance in cognitive task performance (δ) from variance that is unrelated to dementia processes (*g*’); and 3) cognitive domains for verbal ability, episodic memory, executive function and processing speed.

## 2 Methods

### 2.1 Participants

Participants in this study are from the Personality and Total Health (PATH) Through Life Project, which has been described in detail elsewhere [28]. Briefly, participants were randomly sampled from the electoral rolls of the city of Canberra and the neighbouring town of Queanbeyan, Australia, and were recruited into one of three cohorts based on age at baseline, with follow up occurring at 4-year intervals for a total of 12 years. The focus of the present study is on data collected at Wave 4 in the 60+ cohort (age 60-64 at baseline), as this wave contained an expanded cognitive test battery, activities of daily living, and additional questions relating to the ANU-ADRI. Interviews were conducted in 2014-2015 for 1,645 participants (65% retention from baseline). Individuals were excluded if they were not of European ancestry (*n* = 55), had dementia (*n* = 30), MCI (*n* = 144), MMSE ≤ 24 (*n* = 52), missing genetic data (*n* = 168), or had a self-reported history of stroke, transient ischemic attack, epilepsy, brain tumours, or brain infection (*n* = 243). This left a final sample of 1,061. Written informed consent was obtained from all participants. This study was approved by the Human Research Ethics Committee of The Australian National University.

### 2.2 ANU-ADRI

The development of the ANU-ADRI and the methodology underlying its computation have been described previously [25]. Briefly, the ANU-ADRI can be computed based on up to 15 domains, with the present score was comprised of 11 domains including age, sex, alcohol consumption, diabetes, education, depression, traumatic brain injury, smoking, social engagement, cognitive activities and dietary fish intake (see Supplementary Materials). For each domain, points weighted by each risk factors effect size are allocated based on varying levels within the domain and overall composite score computed as the sum of all available sub-scores. Three domains were not included in this analysis, namely BMI and hypercholesterolemia (as increased risk of dementia is associated with midlife rather than late-life) and pesticide exposure as data was not available in PATH. However, the ANU-ADRI is still predictive of dementia and MCI even when a subset of variables is used indicating that the variables used in the construction of the ANU-ADRI for this study is sufficient [24,26]. The ANU-ADRI total score was transformed into a Z-score.

### 2.3 Genotyping

The top-hit late-onset Alzheimer’s disease (LOAD) single nucleotide polymorphisms (SNPs) identified via genome wide association studies [29-34] from 23 loci (Supplementary Table 1) were genotyped as previously described [18]. The two SNPs defining the *APOE* alleles were genotyped separately using Taqman assays previously described [35]. A weighted explained variance genetic risk score (EV-GRS) for LOAD was constructed. The EV-GRS is the sum of all the LOAD risk alleles across the individuals, weighted by the minor allele frequency (MAF) and the odds ratio associated with LOAD. The formula for calculating the EV-GRS is described in the Supplementary Materials. The EV-GRS was transformed into a z-score.

### 2.4 Activities of daily living

Informants (*n* = 1438) nominated by PATH participants were asked to rate participants on deficits in the performance of everyday activities using the Bayer Activities of Daily Living (B-ADL) [36] in a telephone interview. B-ADL is comprised of 25 items, with the first two evaluating participants’ ability to manage everyday activities and taking care of themselves. Items three to twenty assess specific tasks of everyday life, while the remaining five relate to cognitive functions important for performing activities of daily living. Informants were asked to rate participants on a scale of 1 (never) to 10 (always), with an option of not applicable. Individual item scores were summed, with the total divided by the number of items rated with a score, providing a final score of between 1 and 10. The B-ADL was reverse coded so that higher scores indicated better function and transformed into a Z-Score.

### 2.5 Cognitive test battery

The PATH cognitive test battery included measures used at previous waves as well as additional tests that were added to the battery to enable clinical diagnoses according to DSM5 criteria [37]. The test battery assessed four cognitive domains. More detailed descriptions of the individual tests can be found in the Supplementary Materials.

*Episodic Memory* was assessed using the Immediate and Delayed Recall of the first trial of the California Verbal Learning Test (IR and DR) [38] and the Benton Visual Retention Test (BVRT) Administration B [39].

*Verbal Ability/Fluency* was assessed using the Spot-the-Word test [40], Boston Naming Test (BNT) [41] and the Controlled Oral Word Association Test (COWAT) [42].

*Perceptual Speed* was assessed using the Symbol Digit Modalities test (SDMT) [43], the Trail Making Test part A (TMT-A) [42] and Simple and Choice Reaction Time (SRT & CRT) [44].

*Executive Function* was assessed using the Victoria Stroop Test interference score [45], the Zoo Map test from the Behavioural Assessment of the Dysexecutive Syndrome Test Battery [46], the Trail Making Test Part B (TMT-B) [42], and Digit Span Backwards (DSB) from the Wechsler Memory Scale [47], with participants were scored based on the number of correct trials (DSB total score) and the longest sequence repeated backwards (DSB Sequence length).

The means and standard deviations for the raw cognitive tests are presented in Table 1. TMT-A, TMT-B, Stroop, SRT and CRT scores were reversed coded so that a higher score also indicates better performance. For TMT-A, TMT-B and Stroop Interference, the Skew was >± 3 or Kurtosis was >± 10, as such extreme outliers (99% percentile on TMT-A; 99.8% percentile on TMT-B and Stroop) were winsorized so that the cognitive test performance distribution approached normality to facilitate estimation of the CFA models [48]. Cognitive test scores were transformed into Z-scores.

**Table 1:**
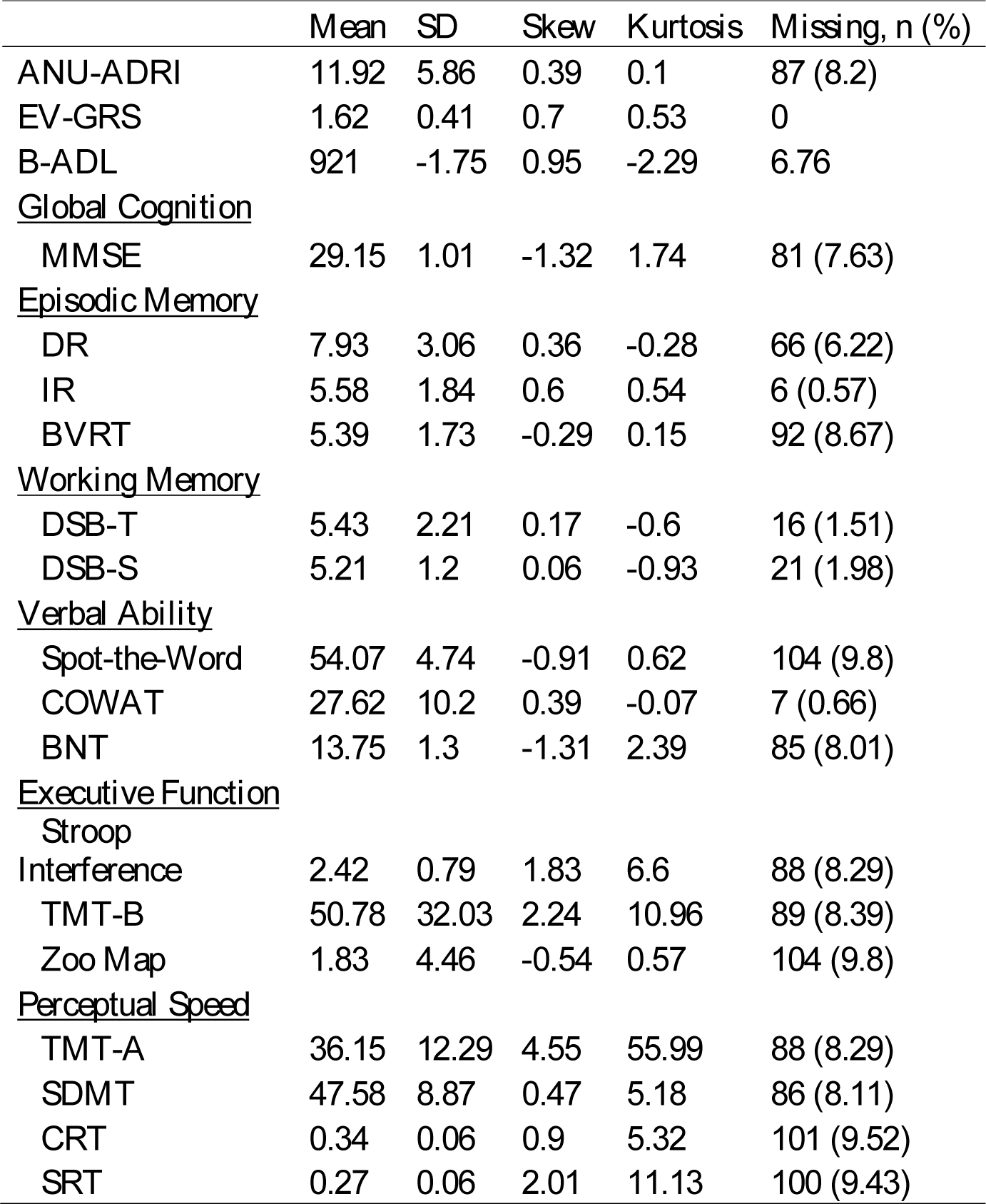
Descriptive statistics of the raw ANU-ADRI, EV-GRS, B-ADL and cognitive test scores available at wave 4 in the PATH 60s Cohort.

*Abbreviations.* ANU-ADRI, Australian National University Alzheimer’s Disease Risk Index; EV-GRS, Explained Variance Genetic Risk Score; B-ADL, Bayer Activities of Daily Living; MMSE, Mini-Mental Stat examination; IR, Immediate Recall of the first trial of the California Verbal Learning Test; DR, Delayed Recall of the first trial of the California Verbal Learning Test; BVRT, Benton Visual Retention Test; Spot-Word, the Spot-the-Word test; COWAT, Controlled Oral Word Association Test; SDMT, Symbol Digit Modalities test; TMT-A, Trail Making Test part A; SRT, Simple Reaction Time; CRT, Choice Reaction Time; Stroop, Victoria Stroop Test interference score; Zoo, Zoo Map test; TMT-B, Trail Making Test Part B; DSB-T, the total number of correct trials in Digit Span Backwards; DSB-S, the longest sequence repeated backwards on the Digit Span Backwards.

### 2.6 Statistical Analysis

All statistical analyses were performed using R version 3.4.2 software [49]. All missing values for the cognitive tests, B-ADL and the ANU-ADRI (see Table 1) were imputed using a Random Forest algorithm from the ‘missForrest’ R package [50].

Confirmatory factor analysis (CFA) can be used to represent the correlations between a number of observed variables (indicators) in terms of a smaller number of unobserved latent variables (factors). CFA is a hypothesis driven approach in which the loading of an indicator onto a factor is based on *a priori* evidence and theory. CFA models were estimated using mean-adjusted weighted least squares (WLSM) in the ‘lavaan’ R package [49,51].

Three separate CFA models were constructed (Figure 1). First, Model 1 (Figure 1a) is a single factor model in which all the individual cognitive tests loaded onto a single latent factor representing general cognitive ability (g) was constructed. Second, Model 2 (Figure 1b), presents bifactor model distinguishing dementia-related variance in cognitive task performance (δ) from variance that is unrelated to dementia processes (*g*’) was constructed. Individual cognitive tests were loaded onto both g’ and δ, while the B-ADL loaded only onto δ. As such, the latent variable δ represents the shared variance between cognitive and functional measures. Third, Model 3 (Figure 1c) is a multi-factor cognitive domain CFA model, which shows that the individual cognitive tests were loaded onto four factors representing the cognitive domains of Verbal Ability (VA), Episodic Memory (EM), Executive Function (EF) and Processing Speed (PS).

**Figure 1:**
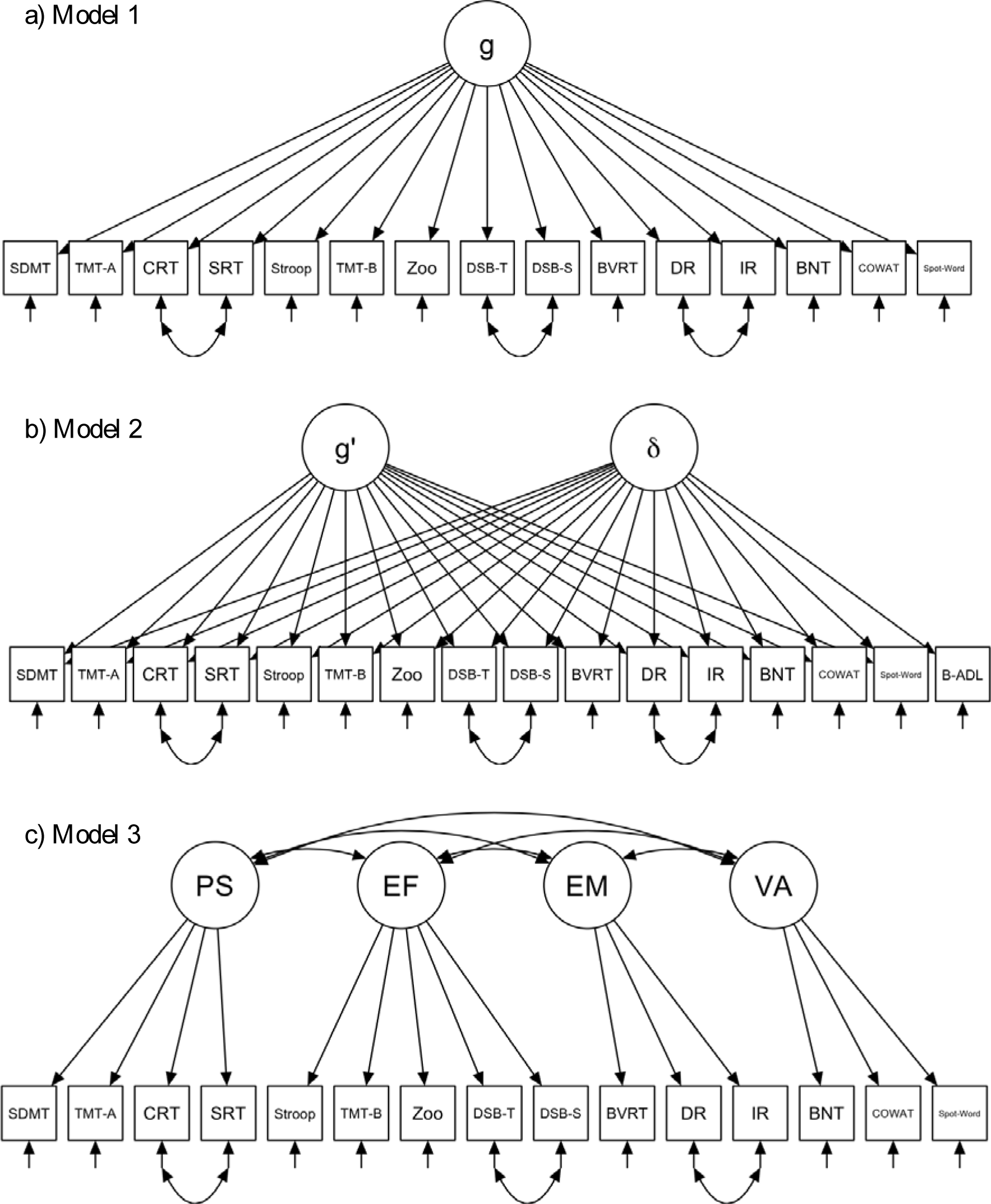
Path diagrams for the hypothesized factor structures for the three confirmatory factor analysis models. a) Model 1: single factor for general intelligence. b) Model 2: bifactor model that distinguishes dementia-related variance in cognitive task performance (δ) from variance that is unrelated to dementia processes (*g*’). c) Model 3: multifactor model composed of four cognitive domains verbal ability, episodic memory, executive function and processing speed. *Abbreviations*. IR, Immediate Recall of the first trial of the California Verbal Learning Test; DR, Delayed Recall of the first trial of the California Verbal Learning Test; BVRT, Benton Visual Retention Test; Spot-Word, the Spot-the-Word test; COWAT, Controlled Oral Word Association Test; SDMT, Symbol Digit Modalities test; TMT-A, Trail Making Test part A; SRT, Simple Reaction Time; CRT, Choice Reaction Time; Stroop, Victoria Stroop Test interference score; Zoo, Zoo Map test; TMT-B, Trail Making Test Part B; DSB-T, the total number of correct trials in Digit Span Backwards; DSB-S, the longest sequence repeated backwards on the Digit Span Backwards.

For Models 1-3, error covariances were included between the IR–DR, DSB-T–DSB-S and SRT–CRT items to account for method effects (where additional covariation among indicators is introduced due the measurement approach). The latent variables factor variances were fixed to 1 to allow for all factor loadings to be estimated. Model fit was evaluated with multiple indices of model fitness including Comparative Fit Index (CFI), Root Mean Square Error of Approximation (RMSEA) and Standardized Root Mean Square Residual (SRMSR) [52]. A CFI > 0.9, RMSEA and SRMSR < 0.08 were considered good estimates of model fit.

Under a structural equation modelling framework, the ANU-ADRI and the EV-GRS were introduced as exogenous indicator variables into Models 1-3 to examine direct effect of the ANU-ADRI and the EV-GRS on latent cognitive factors.

## 3 Results

Distributions for the ANU-ADRI, EV-GRS, B-ADL and the individual cognitive test scores are described in Table 1. The frequencies of the individual sub-indices of the ANU-ADRI are presented in Table 2.

**Table 2:**
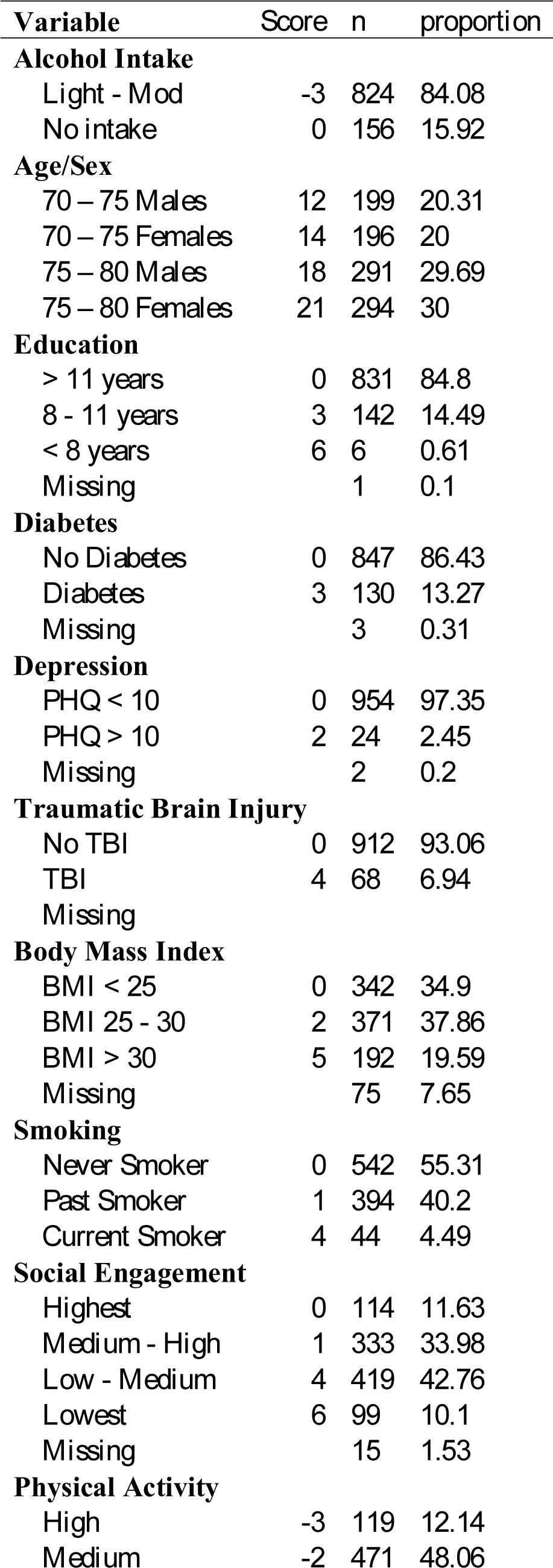
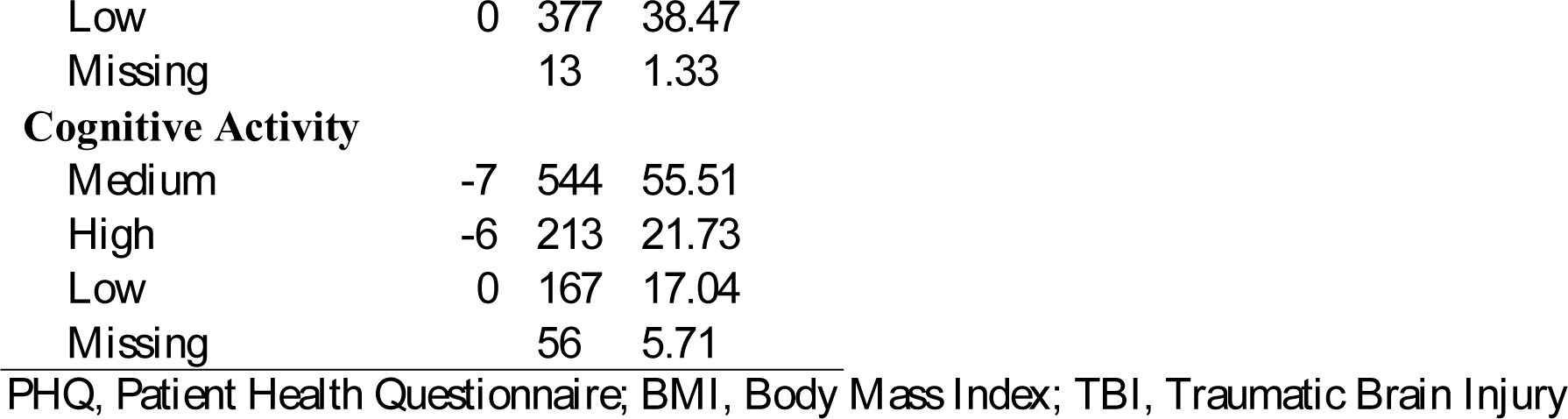
Frequencies of ANU-ADRI Subindices.

### 3.1 Model 1: ‘g’

Fit statistics and standardised parameter estimates for the single-factor ‘g’ CFA model are presented in Table 3 and were acceptable, indicating that the overall fit of Model 1 provided support for the hypothesized structure of the cognitive tests. All the factor loadings, except for SRT and Zoo Map, were above 0.30, ranging in absolute value from 0.33 – 0.64 and thus accounting for between 10.9 – 40.9% of the variance in general cognitive ability.

**Table 3:** Standardized parameter estimates and model fit for Model 1 (‘g’), Model 2 (g’ & δ) model, Model 3 (cognitive domain)

| Indicator/Factor | Model 2<br>Estimate (SE) | Model 2<br>Estimate (SE) | Model 3<br>Estimate (SE) |
| --- | --- | --- | --- |
| <u>Factor Loadings</u> |  |  |  |
| **g** |  |  |  |
| Processing Speed |  |  |  |
| SDMT | 0.64 (0.03)*** | 0.16 (0.08)* | 0.81 (0.04)*** |
| TMT-A | 0.41 (0.03)*** | -0.03 (0.07) | 0.52 (0.03)*** |
| CRT | 0.33 (0.03)*** | 0.06 (0.05) | 0.41 (0.04)*** |
| SRT | 0.28 (0.04)*** | 0.08 (0.05) | 0.34 (0.04)*** |
| Executive Function |  |  |  |
| Stroop | 0.48 (0.03)*** | 0.27 (0.06)*** | 0.51 (0.03)*** |
| TMT-B | 0.57 (0.03)*** | 0.3 (0.06)*** | 0.61 (0.03)*** |
| Zoo Map | 0.29 (0.03)*** | 0.2 (0.05)*** | 0.31 (0.03)*** |
| DSB-T | 0.40 (0.03)*** | 0.37 (0.04)*** | 0.43 (0.03)*** |
| DSB-S | 0.38 (0.03)*** | 0.35 (0.04)*** | 0.41 (0.03)*** |
| Episodic Memory |  |  |  |
| BVRT | 0.55 (0.03)*** | 0.35 (0.06)*** | 0.59 (0.03)*** |
| DR | 0.43 (0.03)*** | 0.29 (0.05)*** | 0.46 (0.03)*** |
| IR | 0.39 (0.03)*** | 0.31 (0.04)*** | 0.42 (0.03)*** |
| Verbal Ability |  |  |  |
| BNT | 0.44 (0.04)*** | 0.36 (0.05)*** | 0.49 (0.04)*** |
| COWAT | 0.59 (0.02)*** | 0.54 (0.05)*** | 0.68 (0.03)*** |
| Spot-the-Word | 0.58 (0.03)*** | 0.67 (0.04)*** | 0.69 (0.03)*** |
| **$\delta$** |  |  |  |
| SDMT | - | 0.72 (0.04)*** | - |
| TMT-A | - | 0.61 (0.04)*** | - |
| CRT | - | 0.4 (0.04)*** | - |
| SRT | - | 0.31 (0.04)*** | - |
| Stroop | - | 0.39 (0.05)*** | - |
| TMT-B | - | 0.5 (0.05)*** | - |
| Zoo Map | - | 0.21 (0.04)*** | - |
| DSB-T | - | 0.21 (0.05)*** | - |
| DSB-S | - | 0.2 (0.05)*** | - |
| BVRT | - | 0.41 (0.05)*** | - |
| DR | - | 0.32 (0.04)*** | - |
| IR | - | 0.24 (0.05)*** | - |
| BNT | - | 0.26 (0.06)*** | - |
| COWAT | - | 0.32 (0.06)*** | - |
| Spot-the-Word | - | 0.24 (0.07)*** | - |
| Bayer ADL | - | 0.43 (0.04)*** | - |
| <u>Variances</u> |  |  |  |
| SDMT | 0.6 (0.04)*** | 0.46 (0.05)*** | 0.35 (0.06)*** |
| TMT-A | 0.83 (0.03)*** | 0.62 (0.05)*** | 0.73 (0.03)*** |
| CRT | 0.89 (0.02)*** | 0.83 (0.03)*** | 0.83 (0.03)*** |
| SRT | 0.92 (0.02)*** | 0.9 (0.02)*** | 0.88 (0.03)*** |
| Stroop | 0.77 (0.03)*** | 0.77 (0.03)*** | 0.74 (0.03)*** |
| TMT-B | 0.67 (0.03)*** | 0.67 (0.03)*** | 0.62 (0.04)*** |
| Zoo Map | 0.91 (0.02)*** | 0.92 (0.02)*** | 0.9 (0.02)*** |
| DSB-T | 0.84 (0.02)*** | 0.82 (0.03)*** | 0.81 (0.03)*** |
| DSB-S | 0.86 (0.02)*** | 0.84 (0.03)*** | 0.83 (0.03)*** |
| BVRT | 0.7 (0.03)*** | 0.71 (0.03)*** | 0.65 (0.04)*** |
| DR | 0.82 (0.02)*** | 0.82 (0.02)*** | 0.79 (0.03)*** |
| IR | 0.85 (0.02)*** | 0.84 (0.02)*** | 0.82 (0.03)*** |
| BNT | 0.81 (0.03)*** | 0.8 (0.03)*** | 0.76 (0.04)*** |
| COWAT | 0.65 (0.03)*** | 0.61 (0.03)*** | 0.54 (0.04)*** |
| Spot-the-Word | 0.66 (0.03)*** | 0.5 (0.04)*** | 0.53 (0.04)*** |
| B-ADL | - | 0.81 (0.04)*** | - |
| <u>Covariances</u> |  |  |  |
| PS – EF | - | - | 0.72 (0.05)*** |
| PS – EM | - | - | 0.69 (0.05)*** |
| PS – VA | - | - | 0.56 (0.04)*** |
| EF – EM | - | - | 0.86 (0.05)*** |
| EF – VA | - | - | 0.74 (0.05)*** |
| EM – VA | - | - | 0.82 (0.05)*** |
| DR – IR | 0.93 (0)*** | 0.92 (0)*** | 0.45 (0.03)*** |
| DSB-T – DSB-S | 0.47 (0.03)*** | 0.47 (0.03)*** | 0.92 (0)*** |
| CRT – SRT | 0.75 (0.02)*** | 0.74 (0.02)*** | 0.73 (0.02)*** |
| <u>Model Fit</u> |  |  |  |
| Model $\chi^2$ (df) | 593.53 (102) | 509.77 (102) | 486 (96) |
| CFI | 0.90 | 0.921 | 0.921 |
| RMSEA (90% CI) | 0.070 (0.066 - 0.074) | 0.065 (0.061 - 0.069) | 0.065 (0.061 - 0.069) |
| SRMR | 0.068 | 0.058 | 0.060 |

\* $p < 0.05$ ; \*\*\* $p < 0.001$

*Abbreviations.* ANU-ADRI, Australian National University Alzheimer’s Disease Risk Index; EV-GRS, Explained Variance Genetic Risk Score; B-ADL, Bayer Activities of Daily Living; MMSE, Mini-Mental Stat examination; IR, Immediate Recall of the first trial of the California Verbal Learning Test; DR, Delayed Recall of the first trial of the California Verbal Learning Test; BVRT, Benton Visual Retention Test; Spot-Word, the Spot-the-Word test; COWAT, Controlled Oral Word Association Test; SDMT, Symbol Digit Modalities test; TMT-A, Trail Making Test part A; SRT, Simple Reaction Time; CRT, Choice Reaction Time; Stroop, Victoria Stroop Test interference score; Zoo, Zoo Map test; TMT-B, Trail Making Test Part B; DSB-T, the total number of correct trials in Digit Span Backwards; DSB-S, the longest sequence repeated backwards on the Digit Span Backwards; VA, Verbal ability; EM, Episodic Memory; EF, Executive Function; PS, Processing Speed.

The ANU-ADRI was significantly associated with general cognitive ability. Specifically, a one SD increase in the ANU-ADRI (5.86 points) corresponded to a decrease of -0.40 (95% CI: -0.37 – -0.43) in ‘g’ and accounted for 16.2% of variance (Table 4; Supplementary Table 2). The association between the EV-GRS and ‘g’ was non-significant (Table 4; Supplementary Table 2).

**Table 4:** Standardized Regression Estimates for the direct effect of the ANU-ADRI and EV-GRS on cognitive performance.

|  | ANU-ADRI<br>Estimate (SE) | EV-GRS<br>Estimate (SE) |
| --- | --- | --- |
| Single-Factor ‘g’ |  |  |
| g | -0.4 (0.03)*** | -0.05 (0.03) |
| Bifactor g’ & δ |  |  |
| g’ | -0.18 (0.06)** | 0.001 (0.04) |
| δ | -0.4 (0.04)*** | -0.08 (0.04)* |
| Multi-factor |  |  |
| Cognitive Domain |  |  |
| PS | -0.40 (0.04)*** | -0.06 (0.04) |
| EF | -0.38 (0.04)*** | -0.02 (0.04) |
| EM | -0.34 (0.04)*** | -0.10 (0.05)* |
| VA | -0.29 (0.04)*** | -0.02 (0.04) |

\* $p < 0.05$ ; \*\*\* $p < 0.001$

*Abbreviations.* ANU-ADRI, Australian National University Alzheimer’s Disease Risk Index; EV-GRS, Explained Variance Genetic Risk Score; VA, Verbal ability; EM, Episodic Memory; EF, Executive Function; PS, Processing Speed.

### 3.2 Model 2: g’ and δ

Standardised parameter estimates, and model fit indices for Model 2 are presented in Table 3. The bifactor structure of Model 2 demonstrated good overall fit with the data. All the cognitive variables and the B-ADL loaded significantly on δ, with all loadings above 0.30. In comparison to Model 1, factor loadings on g’ were reduced and the loading of TMT-A, SRT and CRT on g’ were no longer significant.

The ANU-ADRI was significantly associated with both δ (β = -0.4; 95% CI = -0.586 – -0.367) and g’ (β = -0.18; -0.332 – -0.059) (Table 4; Supplementary Table 2). A one SD increase in the ANU-ADRI was associated with worse performance in both δ and g’. However, a larger decrease in variance explained (16.3%) was observed for δ as compared with to g’ (3.2%). For the EV-GRS, a significant association was observed for δ (β = - 0.08; 95% CI = -0.166 – -0.003), which explained 0.58% of the variation in δ.

### 3.3 Model 3: Cognitive Domains

Fit statistics and standardised parameter estimates for the four-factor cognitive domain model are provided in Table 3. All the model fit indices were acceptable, indicating that the overall fit of the CFA model provided support for the hypothesized cognitive domain structure. The standardized factor loadings confirmed that each of the cognitive domains were well defined by the individual cognitive tests and were all above 0.30, ranging in absolute value from 0.31 – 0.81 and thus accounting for between 9.6 - 65% of the variance. All the cognitive domains were positively inter-correlated, with absolute values ranging from 0.56 (PS – VA) to 0.86 (EF – EM).

A higher ANU-ADRI score was associated with worse performance on all the cognitive factor scores, with a one SD increase in the ANU-ADRI leading to worse cognitive performance ranging from -0.29 (95% CI: -0.36 – -0.22) for VA to -0.40 (95% CI: -0.47 – - 0.33) for PS (Table 4; Supplementary Table 2). The ANU-ADRI accounted for notable factor score variation in PS (16.3%), EF (14.7%), EM (11.8%) and VA (8.5%). A higher EV-GRS was associated with a worse performance in EM, with a one SD increase leading to a -0.098 (95% CI: -0.188 – -0.008) decrease in EM, accounting for 0.96% of the variation in EM (Table 4; Supplementary Table 2).

## 4 Discussion

This study’s main finding was that the ANU-ADRI was cross-sectionally broadly associated with cognitive performance. A higher score was significantly associated with worse performance in ‘g’, representing a latent variable of general cognitive ability. When we further partitioned g into two independent factors, g’ and δ, the ANU-ADRI was associated with worse factor scores for both indicators. Given that deficits in cognition and functional status are key characteristics for a clinical diagnosis of dementia, the latent dementia construct ‘δ’ was conceptualised as a measure to detect early cognitive change and concomitant functional decline associated with neurodegenerative disease [9]. In contrast, g’ reflects cognitive task performance that is unrelated to functional decline caused by neurodegenerative disease [9]. Accordingly, we observed that the effect size of the ANU-ADRI – δ association was larger in comparison to the ANU-ADRI – g’ association. This suggests that while the ANU-ADRI is broadly negatively correlated with cognitive performance, it is more specifically associated with dementia-related processes.

Differences in cognitive ability across domains reflect neuroanatomical differences in localized regional structures/networks and the connectivity of those networks. As such, the differential association of risk and protective factors with specific cognitive domains may reflect associations with particular neuroanatomical structures. The ANU-ADRI was associated with worse performance across all four cognitive domains, however larger effects were observed for processing speed and executive function. Similarly, the CAIDE risk score is broadly associated with poorer cognitive function, with larger deficits in executive functioning and processing speed, in comparison to memory [53]. Deficits in processing speed and executive functioning are characteristic of vascular dementia (VaD) caused by cerebrovascular disease such as infarcts, lacunas, hippocampal sclerosis and white matter lesions [54,55]. Supporting the link between CAIDE and cerebrovascular pathology, a higher baseline score was associated with more severe deep white mater lesions, lower grey matter and hippocampal volume, but not with amyloid accumulation, 20 – 30 years later [53]. This suggests that the ANU-ADRI may also be particularly sensitive to changes in cognitive performance resulting from cerebrovascular disease. The ANU-ADRI has been previously observed to be associated with lower brain volumes in cortical grey matter and the default mode network [27], but associations with cerebrovascular pathology are yet to be evaluated.

In conjunction with the ANU-ADRI, we evaluated the association of an AD GRS with cognitive performance. The GRS, in comparison to the ANU-ADRI, was not significantly associated with general cognitive ability factor in Model 1. In Model 2, a higher EV-GRS was significantly associated only with worse performance in δ. Notably, the EV-GRS effect size (0.58%) was substantially smaller than the ANU-ADRI effect size (16.3%). To our knowledge this is the first study to examine the association of an AD GRS with δ, though *APOE* ε4 has previously been associated with δ [12]. The association of AD genetic markers with δ and not g’ suggests that these risk loci may not promote neural damage independently of AD pathogenesis. As such, these results provide additional support for the validity of δ as a latent dementia phenotype representing dementia severity. In Model 3, a higher EV-GRS was only associated with worse episodic memory performance. Impairment in episodic memory is usually the earliest and most salient characteristic of AD, with deficits in other cognitive domains observed with increasing AD severity [56]. Overall, that the EV-GRS was selectively associated with preclinical memory performance – and that the effect sizes were generally small – likely reflects the fact that the genes comprising the EV-GRS were identified for their associations to AD and its underlying neuropathology.

This study has a number of strengths including a large sample size, a comprehensive cognitive test battery allowing for the modelling of latent cognitive factors, a narrow age range cohort, and the ability to compare an AD environmental/lifestyle risk score to an AD genetic risk score. The main limitation of the current study is its cross-sectional design which has limited ability to evaluate causal relationships and are potentially subject to greater confounding due to cohort effects in comparison to prospective studies. As such, further validation of the ANU-ADRI with cognitive decline is required. Additionally, while PATH was recruited as a representative sample, the educational attainment of the cohort is above the national average and it is a predominantly Caucasian sample, potentially limiting the generalizability of the results of this study.

In conclusion, a higher ANU-ADRI score is associated with worse performance in dementia-related variance in cognitive task performance in comparison to variance in cognitive function unrelated to dementia processes. Additionally, more specific associations were observed with perceptual speed, executive function, episodic memory and verbal ability. In contrast, an AD GRS was specifically associated with dementia-related variance in cognitive task performance and episodic memory. These results provide additional support for using the ANU-ADRI across the cognitive spectrum in individual patient assessment to inform intervention and treatment strategies aimed at delaying dementia.

## Acknowledgments

We thank the participants in the PATH study. We thank the investigators in the PATH study: Peter Butterworth, Andrew Mackinnon, Anthony Jorm, Bryan Rodgers, Helen Christensen, Patricia Jacomb, Karen Mawell and Simon Easteal. The study was supported by the National Health and Medical Research Council (NHMRC) grants 179805 and 1002160 and the NHMRC Dementia Collaborative Research Centre Early Diagnosis and Prevention. SJA is funded by the ARC Centre of Excellence in Population Ageing Research, ARC grant CE1101029. NC is funded by NHMRC Research Fellowship number 12010227. KJA is funded by NHMRC Research Fellowship number 1002560. RAD is funded by the National Institutes of Health (National Institute on Aging, R01 AG008235) and the Canadian Consortium on Neurodegeneration in Aging (with funding from the Canadian Institutes of Health Research and partners).

## References

[1] Barnes DE, Yaffe K. The projected effect of risk factor reduction on Alzheimer’s disease prevalence. Lancet Neurol 2011;10:819–28. doi:10.1016/S1474-4422(11)70072-2.

[2] la Torre de JC. Do We Try Mending Humpty Dumpty or Prevent His Fall? An Alzheimer’s Disease Dilemma. J Alzheimers Dis 2015;46:289–96. doi:10.3233/JAD-150124.

[3] Baker JE, Lim YY, Pietrzak RH, Hassenstab J, Snyder PJ, Masters CL, et al Cognitive impairment and decline in cognitively normal older adults with high amyloid-beta: A meta-analysis. Alzheimers Dement (Amst) 2017;6:108–21.

[4] Hedden T, Oh H, Younger AP, Patel TA. Meta-analysis of amyloid-cognition relations in cognitively normal older adults. Neurology 2013;80:1341–8. doi:10.1212/WNL.0b013e31828ab35d.

[5] Stern Y. Cognitive reserve in ageing and Alzheimer’s disease. Lancet Neurol 2012;11:1006–12. doi:10.1016/S1474-4422(12)70191-6.

[6] Tucker-Drob EM, Salthouse TA. Individual Differences in Cognitive Aging. vol. 132. Oxford, UK: Wiley‐Blackwell; 2011. doi:10.1002/9781444343120.ch9.

[7] Spearman C. “General Intelligence,” objectively determined and measured. The American Journal of Psychology 1904;15:201–92.

[8] Carroll JB. Human cognitive abilities: A survey of factor-analytic studies. Cambridge University Press; 1993.

[9] Royall DR, Palmer RF. Getting Past “g”: testing a new model of dementing processes in persons without dementia. J Neuropsychiatry Clin Neurosci 2012;24:37–46. doi:10.1176/appi.neuropsych.11040078.

[10] Royall DR, Palmer RF, O’Bryant SE. Validation of a latent variable representing the dementing process. J Alzheimers Dis 2012;30:639–49. doi:10.3233/JAD-2012-120055.

[11] Royall DR, Lauterbach EC M.D., Kaufer D M.D., Malloy P Ph.D., Coburn KL Ph.D., Black KJ M.D. The Cognitive Correlates of Functional Status: A Review From the Committee on Research of the American Neuropsychiatric Association. J Neuropsychiatry Clin Neurosci 2007;19:249–65. doi:10.1176/jnp.2007.19.3.249.

[12] Gavett BE, John SE, Gurnani AS, Bussell CA, Saurman JL. The Role of Alzheimer“s and Cerebrovascular Pathology in Mediating the Effects of Age, Race, and Apolipoprotein E Genotype on Dementia Severity in Pathologically-Confirmed Alzheimer”s Disease. J Alzheimers Dis 2016;49:531–45. doi:10.3233/JAD-150252.

[13] Koppara A, Wolfsgruber S, Kleineidam L, Schmidtke K, Frolich L, Kurz A, et al The Latent Dementia Phenotype delta is Associated with Cerebrospinal Fluid Biomarkers of Alzheimer’s Disease and Predicts Conversion to Dementia in Subjects with Mild Cognitive Impairment. J Alzheimers Dis 2016;49:547–60. doi:10.3233/JAD-150257.

[14] Gavett BE, Vudy V, Jeffrey M, John SE, Gurnani AS, Adams JW. The delta latent dementia phenotype in the uniform data set: Cross-validation and extension. Neuropsychology 2015;29:344–52. doi:10.1037/neu0000128.

[15] Palmer RF, Royall DR. Future Dementia Severity is Almost Entirely Explained by the Latent Variable delta’s Intercept and Slope. J Alzheimers Dis 2016;49:521–9. doi:10.3233/JAD-150254.

[16] Royall DR, Palmer RF. δ scores predict mild cognitive impairment and Alzheimer’s disease conversions from nondemented states. Alzheimers Dement (Amst) 2017;6:214–21.

[17] Marden JR, Mayeda ER, Walter S, Vivot A, Tchetgen Tchetgen EJ, Kawachi I, et al Using an Alzheimer Disease Polygenic Risk Score to Predict Memory Decline in Black and White Americans Over 14 Years of Follow-up. Alzheimer Dis Assoc Disord 2016;30:195–202. doi:10.1097/WAD.0000000000000137.

[18] Andrews SJ, Das D, Anstey KJ, Easteal S. Late Onset Alzheimer’s Disease Risk Variants in Cognitive Decline: The PATH Through Life Study. Journal of Alzheimer’s Disease 2017;57:423–36. doi:10.3233/JAD-160774.

[19] Norton S, Matthews FE, Barnes DE, Yaffe K, Brayne C. Potential for primary prevention of Alzheimer’s disease: an analysis of population-based data. Lancet Neurol 2014;13:788–94. doi:10.1016/S1474-4422(14)70136-X.

[20] Livingston G, Sommerlad A, Orgeta V, Costafreda SG, Huntley J, Ames D, et al Dementia prevention, intervention, and care. Lancet 2017. doi:10.1016/S0140-6736(17)31363-6.

[21] Ashby-Mitchell K, Burns R, Shaw J, Anstey KJ. Proportion of dementia in Australia explained by common modifiable risk factors. Alzheimer’s Research & Therapy 2017;9:11. doi:10.1186/s13195-017-0238-x.

[22] Baumgart M, Snyder HM, Carrillo MC, Fazio S, Kim H, Johns H. Summary of the evidence on modifiable risk factors for cognitive decline and dementia: A population-based perspective. Alzheimers Dement 2015;11:718–26. doi:10.1016/j.jalz.2015.05.016.

[23] Kivipelto M, Helkala EL, Hänninen T, Laakso MP, Hallikainen M, Alhainen K, et al Midlife vascular risk factors and late-life mild cognitive impairment: A population-based study. Neurology 2001;56:1683–9.

[24] Anstey KJ, Cherbuin N, Herath PM, Qiu C, Kuller LH, López OL, et al A Self-Report Risk Index to Predict Occurrence of Dementia in Three Independent Cohorts of Older Adults: The ANU-ADRI. PLoS One 2014; 9:e86141. doi:10.1371/journal.pone.0086141.

[25] Anstey KJ, Cherbuin N, Herath PM. Development of a New Method for Assessing Global Risk of Alzheimer’s Disease for Use in Population Health Approaches to Prevention. Prev Sci 2013;14:411–21. doi:10.1007/s11121-012-0313-2.

[26] Andrews SJ, Eramudugolla R, Velez JI, Cherbuin N, Easteal S, Anstey KJ. Validating the role of the Australian National University Alzheimer’s Disease Risk Index (ANU-ADRI) and a genetic risk score in progression to cognitive impairment in a population-based cohort of older adults followed for 12 years. Alzheimer’s Research & Therapy 2017;9:318. doi:10.1186/s13195-017-0240-3.

[27] Cherbuin N, Shaw ME, Walsh E, Sachdev P, Anstey KJ. Validated Alzheimer’s Disease Risk Index (ANU-ADRI) is associated with smaller volumes in the default mode network in the early 60s. Brain Imaging Behav 2017;65:550. doi:10.1007/s11682-017-9789-5.

[28] Anstey KJ, Christensen H, Butterworth P, Easteal S, Mackinnon A, Jacomb T, et al Cohort profile: the PATH through life project. Int J Epidemiol 2012;41:951–60. doi:10.1093/ije/dyr025.

[29] Seshadri S, Fitzpatrick AL, Ikram MA, DeStefano AL, Gudnason V, Boada M, et al Genome-wide analysis of genetic loci associated with Alzheimer disease. Jama 2010;303:1832–40. doi:10.1001/jama.2010.574.

[30] Naj AC, Jun G, Beecham GW, Wang L-S, Vardarajan BN, Buros J, et al Common variants at MS4A4/MS4A6E, CD2AP, CD33 and EPHA1 are associated with late-onset Alzheimer’s disease. Nat Genet 2011;43:436–41. doi:10.1038/ng.801.

[31] Hollingworth P, Harold D, Sims R, Gerrish A, Lambert J-C, Carrasquillo MM, et al Common variants at ABCA7, MS4A6A/MS4A4E, EPHA1, CD33 and CD2AP are associated with Alzheimer’s disease. Nat Genet 2011;43:429–35. doi:10.1038/ng.803.

[32] Harold D, Abraham R, Hollingworth P, Sims R, Gerrish A, Hamshere ML, et al Genome-wide association study identifies variants at CLU and PICALM associated with Alzheimer’s disease. Nat Genet 2009;41:1088–93. doi:10.1038/ng.440.

[33] Lambert J-C, Heath S, Even G, Campion D, Sleegers K, Hiltunen M, et al Genome-wide association study identifies variants at CLU and CR1 associated with Alzheimer’s disease. Nat Genet 2009;41:1094–9. doi:10.1038/ng.439.

[34] Lambert JC, Ibrahim-Verbaas CA, Harold D, Naj AC, Sims R, Bellenguez C, et al Meta-analysis of 74,046 individuals identifies 11 new susceptibility loci for Alzheimer’s disease. Nat Genet 2013;45:1452–8. doi:10.1038/ng.2802.

[35] Jorm AF, Mather KA, Butterworth P, Anstey KJ, Christensen H, Easteal S. APOE genotype and cognitive functioning in a large age-stratified population sample. Neuropsychology 2007;21:1–8. doi:10.1037/0894-4105.21.1.1.

[36] Hindmarch I, Lehfeld H, Jongh P, Erzigkeit H. The Bayer Activities of Daily Living Scale (B-ADL). Dement Geriatr Cogn Disord 1998;9.

[37] Association AP. Diagnostic and statistical manual of mental disorders (DSM-5®). American Psychiatric Pub; 2013.

[38] Delis DC, Kramer JH, Kaplan E, Ober BA. California Verbal Learning Test. San Antonio: Psychological Corporation; 1987. doi:10.2307/25678059.

[39] Benton AL. The revised visual retention test: clinical and experimental applications. Psychological Corporation; 1963.

[40] Baddeley A, Emslie H, Nimmo-Smith I. The Spot-the-Word test: a robust estimate of verbal intelligence based on lexical decision. Br J Clin Psychol 1993;32:55–65.

[41] Mack WJ, Freed DM, Williams BW, Henderson VW. Boston Naming Test: shortened versions for use in Alzheimer’s disease. J Gerontol 1992;47:P154–8.

[42] Reitan RM, Wolfson D. The Halstead-Reitan neuropsychological test battery: Theory and clinical interpretation. vol. 4. Reitan Neuropsychology; 1985.

[43] Smith A. Symbol digit modalities test (SDMT) manual (revised) Western Psychological Services. Los Angeles 1982.

[44] Anstey KJ, Dear K, Christensen H, Jorm AF. Biomarkers, health, lifestyle, and demographic variables as correlates of reaction time performance in early, middle, and late adulthood. Q J Exp Psychol A 2005;58:5–21. doi:10.1080/02724980443000232.

[45] Strauss E, Sherman EM, Spreen O. A compendium of neuropsychological tests: Administration, norms, and commentary. American Chemical Society; 2006.

[46] Wilson BA, Evans JJ, Alderman N, Burgess PW, Emslie H. Behavioural assessment of the dysexecutive syndrome. Methodology of Frontal and Executive Function 1997:239–50.

[47] Wechsler D. A standardized memory scale for clinical use. J Psychol 1945;19:87–95.

[48] Harrington D. Confirmatory Factor Analysis 2008.

[49] Ihaka R, Gentleman R. R: a language for data analysis and graphics. J Comput Graph Stat 1996;5:299–314.

[50] Stekhoven DJ, Bühlmann P. MissForest--non-parametric missing value imputation for mixed-type data. Bioinformatics 2012;28:112–8. doi:10.1093/bioinformatics/btr597.

[51] Rosseel Y. lavaan: An R Package for Structural Equation Modeling. J Stat Soft 2012;48:36. doi:10.18637/jss.v048.i02.

[52] Brown TA. Confirmatory Factor Analysis for Applied Research, Second Edition 2014.

[53] Stephen R, Liu Y, Ngandu T, Rinne JO, Kemppainen N, Parkkola R, et al Associations of CAIDE Dementia Risk Score with MRI, PIB-PET measures, and cognition. Journal of Alzheimer’s Disease 2017;59:695–705.

[54] Smits LL, van Harten AC, Pijnenburg YAL, Koedam ELGE, Bouwman FH, Sistermans N, et al Trajectories of cognitive decline in different types of dementia. Psychol Med 2015;45:1051–9. doi:10.1017/S0033291714002153.

[55] John SE, Gurnani AS, Bussell C, Saurman JL, Griffin JW, Gavett BE. The effectiveness and unique contribution of neuropsychological tests and the delta latent phenotype in the differential diagnosis of dementia in the uniform data set. Neuropsychology 2016;30:946–60. doi:10.1037/neu0000315.

[56] Salmon DP, Bondi MW. Neuropsychological Assessment of Dementia. Annual Review of Psychology 2009;60:257–82.

